# Cryo-EM structure of the polycystin 2-l1 ion channel

**DOI:** 10.1101/287367

**Authors:** Raymond E. Hulse, Zongli Li, Rick K. Huang, Jin Zhang, David E. Clapham

## Abstract

We report the near atomic resolution (3.1 Å) of the polycystic kidney disease 2-like 1 (polycystin 2-l1) ion channel. Encoded by PKD2L1, polycystin 2-l1 is a calcium and monovalent cation-permeant ion channel in primary cilia and plasma membranes. The related primary cilium-specific polycystin-2 protein, encoded by PKD2, shares a high degree of sequence similarity, yet has distinct permeability characteristics. Here we show that these differences are reflected in the architecture of polycystin 2-l1.

## Introduction

Revolutionary improvements in resolving protein and cellular structures, and genetic identification of ciliopathies, have created renewed interest in primary cilia. These small protuberances (5-10μm in length) are found in nearly every cell type. Primary cilia house key downstream elements of the sonic hedgehog pathway, which regulates embryonic development and some cancers. Recently, new tools have enabled measurements of other signaling elements within cilia. Primary cilia have elevated resting internal Ca^2+^ concentrations ([Ca^2+^] = 300-700 nM(Delling et al., 2013)), and possess at least two TRP channels, polycystin-2 and polycystin 2-l1. These ion channels are encoded by the PKD2 and PKD2L1 genes of the TRPP subfamily, respectively. Electrophysiological measurements demonstrate that polycystin 2-l1 and polycystin-2 both underlie ionic currents in primary cilia(DeCaen et al., 2013, DeCaen et al., 2016, Kleene and Kleene, 2017). This suggests that voltage gradients and Ca^2+^, in addition to cAMP, are relevant signals within primary cilia.

PKD2 and PKD2L1 share high degrees of sequence identity (52%) and similarity (71%). Mutations in PKD2 account for ~15% of cases of individuals afflicted with Autosomal Dominant Polycystic Kidney Disease (ADPKD). In contrast, there are no diseases currently linked to PKD2L1. Deletion of PKD2 is lethal in mice(Wu et al., 1998), but deletion of PKD2L1 results in occasional gut malrotation, a relatively mild *situs inversus* phenotype(Delling et al., 2013). Finally, while the functional characterization of polycystin 2-l1 protein has been enabled by its plasma membrane expression(DeCaen et al., 2013, DeCaen et al., 2016), polycystin-2 protein function has been hampered by its restriction to surface expression in primary cilia. Recent approaches have been successful, however, in establishing the key biophysical properties of polycystin-2. These approaches include a gain-of-function mutation that enables expression to the plasma membrane in *Xenopus* oocytes(Pavel et al., 2016) and direct patch clamp recording of polycystin-2 in primary cilia(Kleene and Kleene, 2017) (Liu et al., 2018). These studies establish polycystin-2 as a monovalent-selective cation channel with little or no calcium permeation. In contrast, polycystin 2-l1 shows significant and relevant Ca^2+^ conduction(DeCaen et al., 2013, DeCaen et al., 2016). Despite this knowledge, the physiological stimulus for activation and gating is unclear for both channels.

Here we present a structure of the full-length human polycystin 2-l1 protein at 3.1Å resolution using single particle electron cryo-microscopy (cryo-EM). The overall architecture conforms to that of other TRP channels, and in particular to the TRPP polycystin-2 structure. We establish the core structure and point out differences in regions and residues that may account for polycystin-2 and polycystin 2-l1’s distinct permeation and gating.

## Results

We first purified full-length recombinant polycystin 2-l1 protein tagged with maltose binding protein for cryo-EM (Supplemental Figure 1). To determine whether the MBP-tagged protein was functional, we measured single channel activity in reconstituted liposomes under voltage clamp. The channel’s conductance in the same buffer used for purification (HKN, Methods) was 105 pS, consistent with previous single-channel studies from cells expressing polycystin 2-l1(DeCaen et al., 2013) (Supplemental Figure 1D).

Polycystin 2-l1’s structure exhibits many of the hallmarks of TRP channels; it forms a homotetramer with a domain-swapped Voltage Sensing-Like Domain (VSLD, the S1-S4 transmembrane domains) (***Figure 1A***) and shares remarkable architectural similarity to polycystin-2 (global RMSD of 1.7Å measured with polycystin-2 model pdb5T4D). As characterizes group 2 TRP channels (TRPPs and TRPMLs), polycystin 2-l1 has a long S1-S2 extracellular loop, termed the polycystin mucolipin domain (PMD). This domain, a series of 3 *α*-helices and 5 ß-sheets, forms a cover, or lid, above the channel (***Figure 1B***). In contrast to the 3 or 4 observed glycans in polycystin-2 PMDs(Grieben et al., 2017, Shen et al., 2016, Wilkes et al., 2017), polycystin 2-l1’s PMD has one glycan located at residue N207. Notably, the PMD interacts with underlying elements of the pore, the voltage sensing-like domains, and adjacent PMDs.

**Figure 1.**
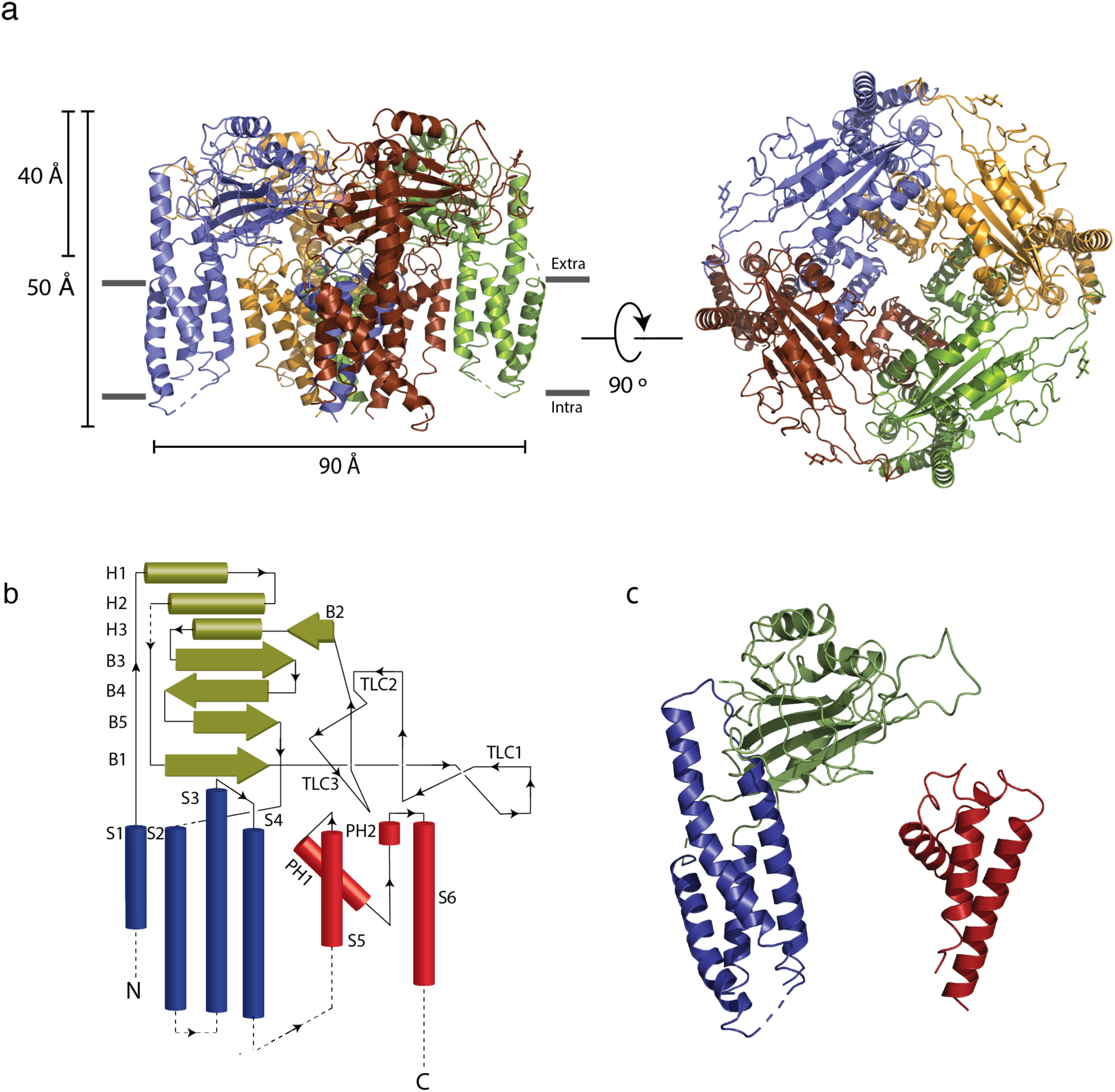
Architecture of polycystin 2-l1. (a) Side view parallel to the membrane and from the top (extracellular) surface. (b) 2D topological representation of the polycystin 2-l1 monomer; voltage sensor-like domain (VSLD, blue), polycystin mucolipin domain (PMD, green), and pore domain (PD, red). (c) Monomer color coded from N terminus (blue) to C terminus (red).

**Figure 2.**
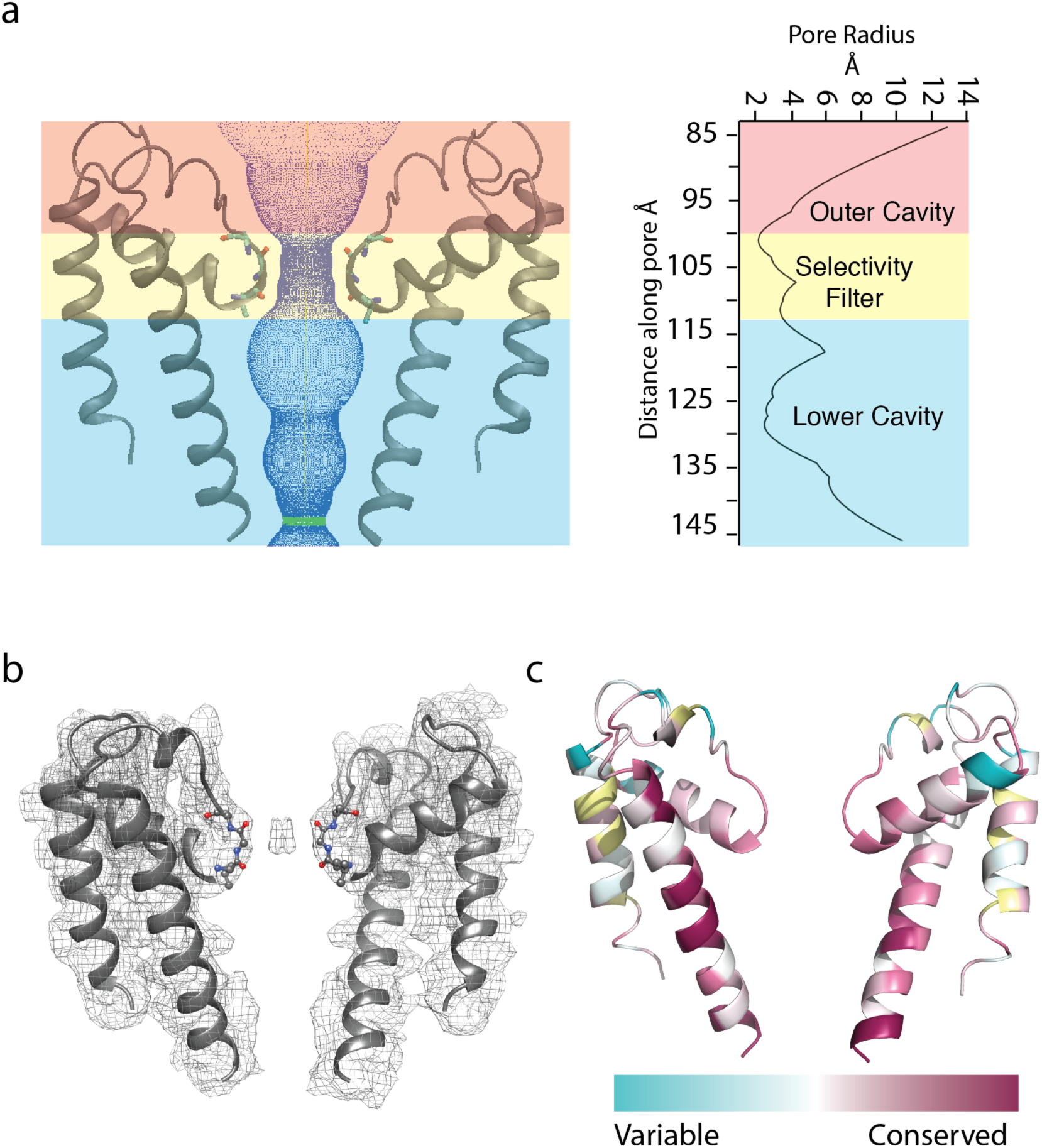
The polycystin 2-l1 pore domain. (a) Path of permeation calculated with HOLE (left). Pore radii along the ion conduction pathway (right). (b) Ion density map superimposed on the polycystin 2-l1 model; contour level 6.0. (c) TRPP family sequence conservation projected onto the polycystin 2-l1 pore domain.

The intracellular face of polycystin 2-l1 is too poorly resolved to permit the N terminus, and the S2-S3 and S4-S5 linker densities to be modeled. Also, despite expression of full-length protein (Supplemental Figure 1) the C terminus of polycystin 2-l1 was not resolved. We conclude that these elements are either unstructured or connected with flexible regions, which, along with the lack of cytoplasmic elements in 2D classification(Shen et al., 2016), prevents model building of these regions.

### The Voltage Sensing-Like Domain

The VSLD is comprised of four transmembrane helical spanning elements that, while similar in structure to the voltage sensing domains of voltage-gated ion channels, do not convert the energy of the transmembrane electric field into pore gating. Indeed, none of the group II TRP channels have significant voltage dependence. Comparison of the VSLD of the two proteins reveals that polycystin 2-l1’s S2 is tilted an additional 9.7° away from the core. Polycystin 2-l1’s S3 is a near-continuous helix extending from the membrane-spanning region into a pocket created by the PMD (***Figure 1c, 3b***). This S3 helix exhibits greater secondary structure than the presumed closed polycystin-2 structure of Shen et al.(Shen et al., 2016), but is similar to that of the multiple- and single-ion models of Wilkes *et al*(Wilkes et al., 2017). The S3 extended helix abuts the PMD of the same monomer but does not display the same cation-π interactions of residues F545 and R320 of polycystin-2. The S3-S4 linker and the top portion of the S4 helix all fit into a cleft of the PMD. In polycystin-2, the S4 adopts a 3_10_-helix configuration (I571 – F579. The density of the S4 helix of polycystin 2-l1 is incomplete at this region so a 3_10_ configuration is not observed. Similarly, the equivalent residues of polycystin-2 that are thought to form a salt-bridge between the S3 and S4 helix (K572, K575 and D511)(Shen et al., 2016), do not have sufficient side chain density to model accurately.

### The Polycystin Mucolipin Domain

The polycystin mucolipin domain (PMD) rests on top of the VSLD and pore domain, with a series of three *α*-helices facing inward towards the funnel/turret of the pore. A series of ß sheets is sandwiched between the *α* helices and a series of loops that interface the extracellular space and the adjacent PMD (***Figure 1b***). A salt-bridge between D295 and K204 near the ß1-ß3 region of the PMD is not observed in polycystin-2 (***Figure 3d***). Finally, a disulfide bond is present between residues C210-C223 in the PMD, similar to several polycystin-2 structures (***Figure 3d***). This element is proposed to stabilize a loop in polycystin-2(Shen et al., 2016, Wilkes et al., 2017) and seems likely to play the same role.

**Figure 3.**
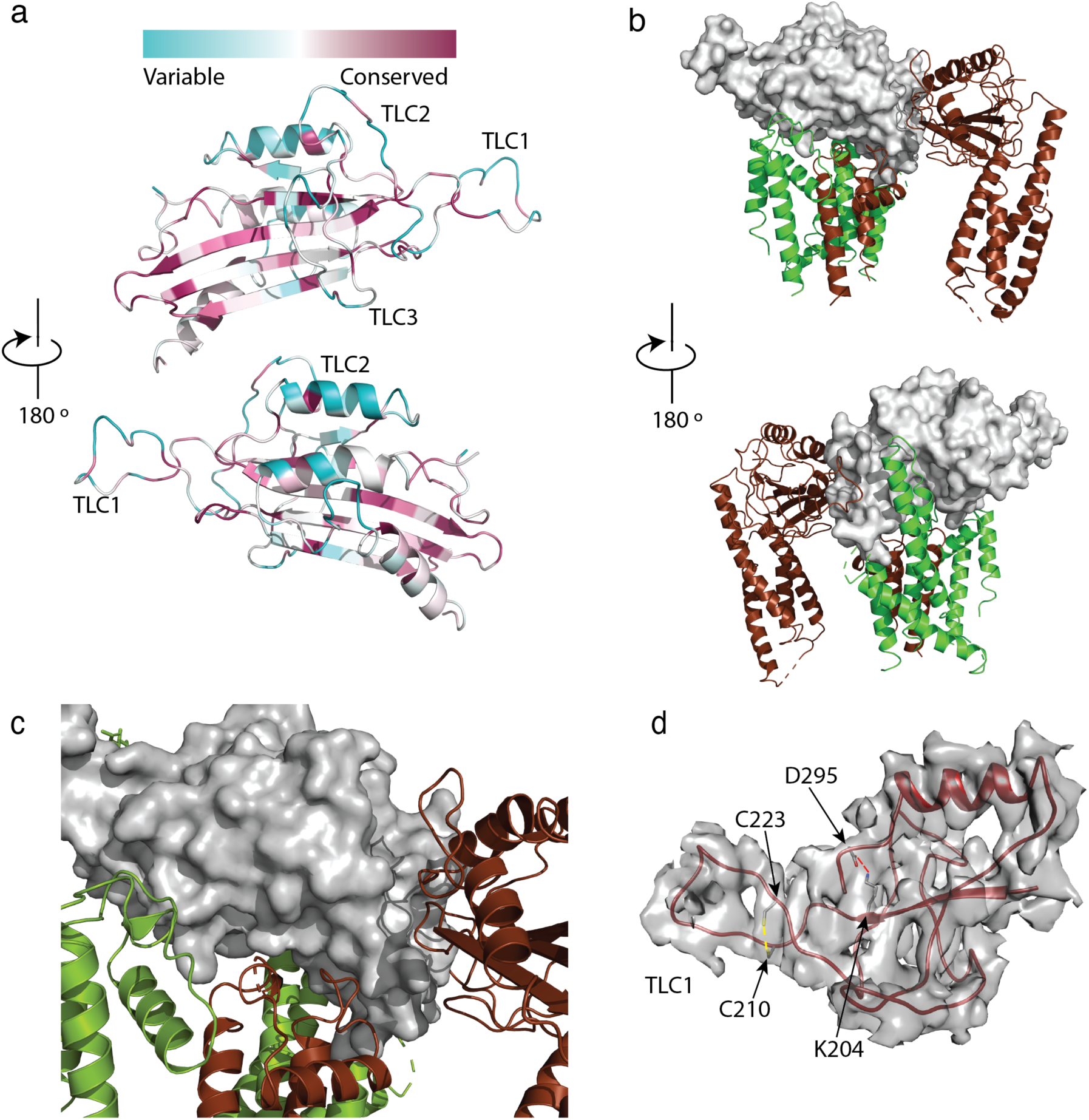
Structure and conservation of the polycystin mucolipin domain, PMD of polycystin 2-l1. – (a) sequence conservation projected onto the PMD domain for polycystin 2-l1 homologs; Three-leaf clover (TLC) domains labeled. (b) The S3, S3-S4 linker, and S4 helix (green) project into a pocket of the PMD (gray) front (top) and back (bottom). (c) Interactions of the PMD (gray) with the pore domain. (d) Disulfide C210-C223 in the TLC1 loop (3.4 Å) and a salt-bridge (red-dash) between D295 and K204 (3.2Å) of the PMD.

### Three leaf clovers of the polycystin mucolipin domain

Grieben *et al.*,(Grieben et al., 2017) described a three-lobed area of the PMD they name a “three-leafed clover” (TLC). Polycystin 2-l1 lacks the small *α* helix in TLC1 of polycystin-2. Interestingly, this region (D208-D225) displays moderately lower sequence conservation among the TRPPs (***Figure 3a***). As noted, TLC1 in polycystin 2 appears to extend into the adjacent PMD domain and interact with the S3 helix and S3-S4 linker(Grieben et al., 2017, Wilkes et al., 2017)). In polycystin 2-l1, the analogous TLC extends from one monomer into the PMD of the adjacent subunit (***Figure 3b***). However, the nature of the interaction differs in that F216 of TLC1 is near W308 of the adjacent subunit’s PMD, representing a possible pi-pi stacking interaction or a hydrophobic pocket (***Figure 4a***). Additionally, N311’s carboxamide oxygen group, located on a hairpin between ß3 and ß4 of the PMD, is within hydrogen bonding distance (3.2Å) of Y224’s amide in TLC1 of the adjacent subunit of polycystin 2-l1 (***Figure 4b***).

**Figure 4.**
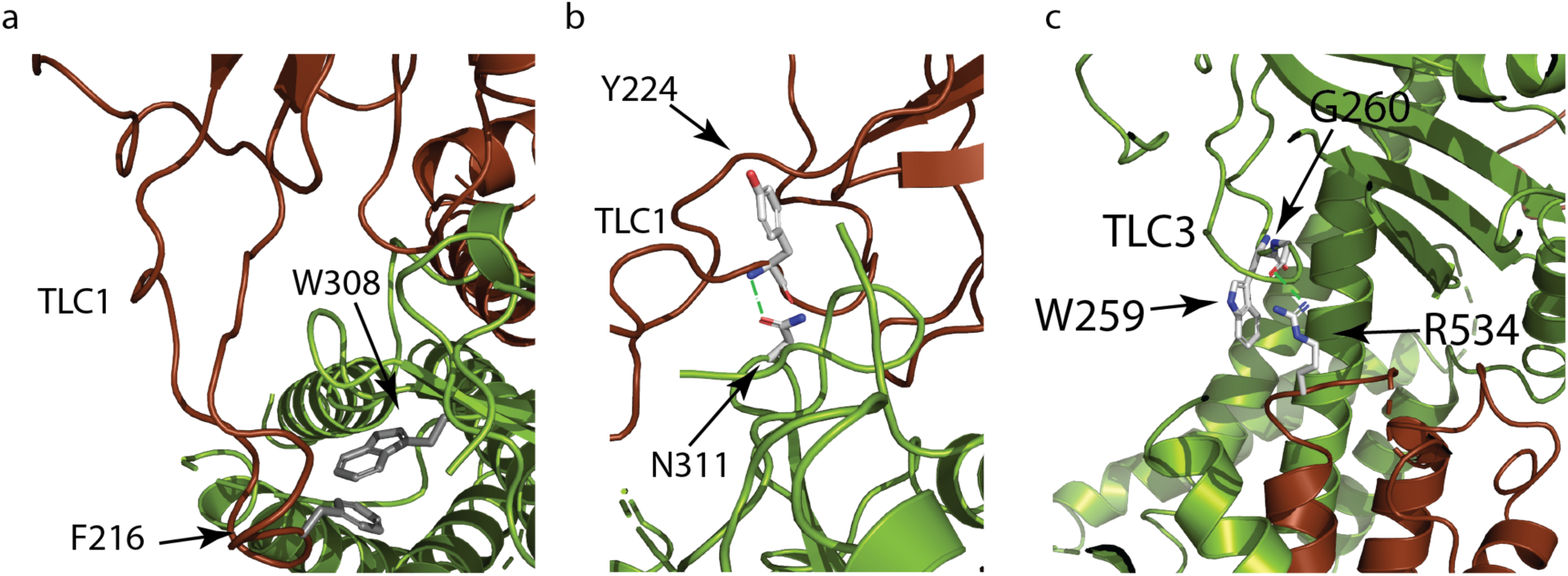
Polycystin mucolipid domain interactions of polycystin 2-l1. – (a) interaction of residue residing in the TLC1 of PMD (F216) with the neighboring PMD (W308). (b) hydrogen bonding interaction (green dash) of the PMD residue N311 with the neighboring PMD residue Y224’s amide (blue). (c)Residue W259 in the TLC3 of PMD forms a cation-π stacking interaction with the upper pore domain residue R534. The guanidino group of R534 also forms a hydrogen bond (green dash) with carbonyl of G260 of the adjacent PMD domain.

### Fenestrations and TLC3

Despite moderately low identity and an increase in charged residues in polycystin 2-l1’s TLC3 (SP<u>DKED</u> versus SVSS<u>ED</u> in polycystin-2), the loop appears essentially the same as in polycystin-2. This element appears to reach under the PMD’s ß sheet core. Proximity of the returning loop of TLC1 towards TLC2, as well as the S5-PH1 loop of the monomer and the PH2-S6 loop adjacent monomer creates four lateral openings at the base of the PMD in polycystin 2-l1, as in polycystin-2. These could present an alternative route for ion permeation(Grieben et al., 2017, Wilkes et al., 2017). This area is less conserved in the polycystin 2-l1 homologs.

Polycystin 2-l1’s PMD also interacts with the adjacent subunit’s upper pore domain. In the polycystin-2 multiple-(pdb5MKF) and single-ion (pdb5MKE) structures, Wilkes *et al*.,(Wilkes et al., 2017) observed glycosylation-dependent interactions of the PMD of one subunit to the loop pore helices of the adjacent(Wilkes et al., 2017), with mutually exclusive glycosylation states. One last interaction between TLC3 and the loop between pore-helix 2 and S6, noted in the polycystin-2 structure(Shen et al., 2016), is seen in polycystin 2-l1: W259, from TLC3, forms a cation-π stacking interaction with R534 and a hydrogen bond with G260’s carbonyl (2.8Å) (***Figure 4c***). This difference may help rationalize the intermediate diameter observed in the polycystin-2 multiple-ion structure (1.4Å)(Wilkes et al., 2017) and the single-ion structure (1.0Å)(Wilkes et al., 2017) where such interactions do not exist, and our polycystin 2-l1 structure (3.1Å).

### The Pore Domain

As in polycystin-2, polycystin 2-l1’s selectivity filter is flanked by two pore-helices, which are themselves flanked by a transmembrane spanning helix. The key filter residues are the residues L521, G522 and D523, in which the carbonyls of L521 and G522 point to the central pore axis (***Figure 2b***). However, D523’s side chain is not well resolved, and the map potential does not support an unambiguous assignment. The pore domain shows moderately strong conservation in S6, and elements of the selectivity filter, as well as pore helix 1 (***Figure 2c***). Notably, one element, K511 in polycystin 2-l1, is highly variable among TRPPs. The corresponding residue is a negatively-charged glutamate in polycystin-2 and an asparagine in PC2L2. The effect of this residue is to confer a net positive electrostatic potential (***Figure 5a***) relative to polycystin-2’s net negative charge.

**Figure 5.**
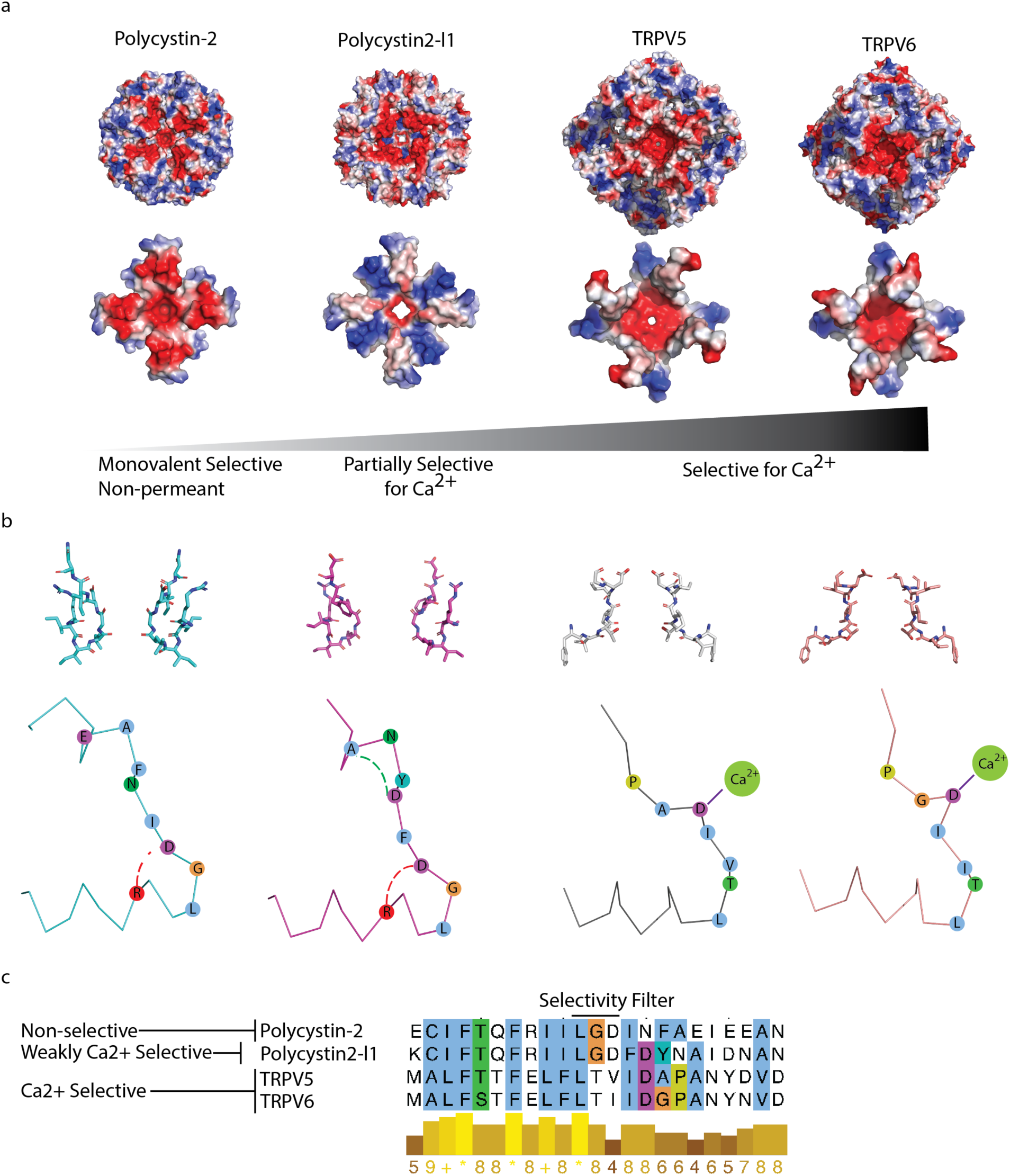
Comparison of charge landscape and local bonding between TRPP and the calcium-selective TRPV5 and TRPV6 channels. a) The extracellular facing (top) and selectivity filter region (bottom) of channels with progressively increasing relative permeability to Ca^2+^. b) Stick representation of the selectivity filter region (top) and simplified diagram (below). Local bonding characteristics are highlighted: conserved salt-bridge (red) interaction of polycystin-2 and polycystin 2-l1, the unique hydrogen bond of polycystin 2-l1 (green), and the direct side-chain coordination of Ca^2+^ in TRPV5 and TRP6. c) Alignment with conservation scoring and clustalW coloring scheme of the selectivity filter region show in b).

Relative to polycystin-2’s 5K47’s pore helix 1, the angle between the beginning of pore helix 1 and the top of the selectivity filter is similar (77° for polycystin 2-l1 and 75° for polycystin-2) compared to the 10° pitch difference between polycystin-2 and TRPV1(Grieben et al., 2017). However, polycystin 2-l1’s pore helix is ~1Å shorter than that of polycystin-2 (measured from beginning to end of the *α*-helix at the C*α*) and so creates a slightly wider filter diameter of 2.1 Å. A second stricture is found in a similar position as the polycystin-2 structures, with a diameter of 2.5Å at N561. The S5 helix appears truncated in the polycystin 2-l1 model due to lack of resolution in this region.

The narrowest distance in the selectivity filter, measured by HOLE(Smart et al., 1996), is 2.1 Å (***Figure 2a***). Set by G522, this represents a larger opening than that in any of the three polycystin-2 structures (1.0Å to 1.4Å)(Grieben et al., 2017, Shen et al., 2016, Wilkes et al., 2017). Such a diameter is sufficient to accommodate a partially hydrated Ca^2+^ ion with an average ~ 2.4 Å metal-oxygen distance(Katz et al., 1996, Marcus, 1988). The observed density in the selectivity filter appears between the two carbonyls of L521 and G522, contrasting with polycystin-2 densities that localize above and below the selectivity filter(Wilkes et al., 2017) (***Figure 2b***). However, cryo-EM cannot determine an ion’s identity with the same degree of reliability enabled by crystallographic anomalous diffraction. Furthermore, although defined buffers are used, we cannot rule out the possibility of less abundant ions occupying the site.

Measured from side-chain to side-chain, polycystin 2-l1’s pore is narrowest at the diagonal from the carbonyl oxygen of G522 to G522 (8.7Å) and becomes slightly larger at G522’s C*_α_* (9.1Å). The pore widens at the subsequent glycine amide bond to 11.0Å, narrows to 9.0Å at the L521 carbonyl, and finally widens to 9.0Å at the C*_α_* of L521. The narrowest aspect of the pore, between the internal-facing carbonyl group of G522 to the C*a* of L521, is not large enough to accommodate a fully hydrated Ca^2+^ ion, as seen in the CaVAb structure (10-12Å)(Tang et al., 2014). Although we do not know whether our polycystin 2-l1 channel structure is in an open, closed, or intermediate state, we conjecture that when a Ca^2+^ does permeate, it must be at least partially dehydrated.

### Ion permeation differences between polycystin-2 and polycystin 2-l1

Polycystin-2 and polycystin 2-l1 core selectivity filters residues (LGD) are conserved, leading us to examine other explanations as to why polycystin 2-l1 conducts Ca^2+^ while polycystin-2 does not(DeCaen et al., 2013, DeCaen et al., 2016, Kleene and Kleene, 2017, Liu et al., 2018). If the pore helix 1-selectivity filter-pore helix 2 region of polycystin 2-l1 (K511-P538) is substituted by polycystin-2’s analogous region (E631-P658), the polycystin 2-l1 chimera has roughly (within 4-fold) similar permeability to Na^+^, K^+^, and Ca^2+^ (Shen et al., 2016). Point mutation experiments in which polycystin 2-l1’s D523 and D525 were mutated to alanine or serine yielded no measurable currents(DeCaen et al., 2013). However, polycystin 2-l1(LG**D**_523_: D523N) was 18-fold, and polycystin 2-l1(LGDF**D**_525_: D525N) 9-fold less calcium-permeant (P_Ca_^2+^/P_Cs_^+^) than wt polycystin 2-l1. Also, and significantly, the D525N mutation reduced outward current block by Ca^2+^, while D523N did not(DeCaen et al., 2016). These experiments indicate that order and placement of negative charge in the narrow region of the pore is important for both Ca^2+^ permeation and block. Higher resolution Cryo-EM structures that identify Ca^2+^ and/or monovalents that are occupying the pore with high certainty (or equivalent crystallographic studies like those of CaVAb(Tang et al., 2014)), and realistic molecular dynamic simulations, should shed light on these issues.

As a final attempt to understand the differences in the relative selectivity of TRP channels, we compared the charge landscape of the pore helix-selectivity filter-pore helix assembly of polycystin-2 and polycystin 2-l1. When the PMD is stripped away and only the pore helices and selectivity filter elements of both channels are visualized, a striking difference appears (**Figure 5a**). Polycystin 2-l1 has a more constrained negative charge density immediately surrounding the pore than does polycystin-2, and its pore-helices form a Maltese cross of net positive charge, rotated 45° to a negative charge distribution cross in polycystin-2. We can compare these charge distributions to the much more Ca^2+^-selective TRPV5 and TRPV6 (P_Ca_ /P_Na_ > 100)(Ramsey et al., 2006) structures(Hughes et al., 2018, McGoldrick et al., 2018) in which the TRPV5/6 channels’ negative charge is much more localized around their respective pores.

## Discussion

Although polycystin 2-l1 and polycystin-2 structures are very similar, there are notable differences that may explain reported functional differences in their selectivity and permeation. For discussion, we divide polycystin 2-l1 into three regions to highlight these differences; the voltage sensing-like domain, the pore mucolipin domain, and the pore. In the voltage sensing-like domain, polycystin 2-l1’s S3 helix and S3-S4 linker extend farther into the PMD (as an *α* helix) than in polycystin-2. In addition, the top of the S3-S4 helical region abuts the pore mucolipin domain. Comparing the pore mucolipin domains, polycystin 2-l1 lacks oligosaccharides that act as bridges between adjacent subunits. Additionally, a salt bridge links D295 and K204 of the TLC1 loop. Finally, although similar, polycystin 2-l1’s pore diameter is slightly larger.

Due to our inchoate understanding of the TRPP family, we must be cautious in linking the observed differences in structure to functional differences. The two main features we seek to understand are selectivity and gating. Comparing selectivities of the two channels, we know that polycystin 2-l1 conducts both monovalent ions and Ca^2+^, while polycystin-2 is monovalent-selective(Kleene and Kleene, 2017, Liu et al., 2018). We explain this difference with two features, pore diameter and electrostatic fields. The relatively larger diameter of polycystin 2-l1’s pore (0.6Å - 1.1Å) may enable partially hydrated calcium’s transit, thus avoiding the large energy required for pore residues to dehydrate calcium. Most striking are the unique electrostatic maps (**Figure 5a**) surrounding the polycystin-2 and polycystin 2-l1 pores. Such differences in charge suggest that the putative energy landscapes create small variations in selectivity, a characteristic suggested for the prokaryotic NaK ion channel(Alam and Jiang, 2011). These slight charge differences, and so landscapes, may impart the subtle relative permeability differences observed among most TRP channels. For organelles in which the TRPP (cilia) and TRPML (endolysosomes) appear, these subtle differences may have driven the evolution of small changes in relative permeation in order to titrate levels of ions within these sub-femtoliter compartments. Research into the relative permeabilities of the related TRPML family of ion channels will further our understanding of mechanisms beyond direct side chain coordination.

TRPPs are not appreciably voltage-gated and polycystin 2-l1 is not mechanically gated(DeCaen et al., 2013), which raises the question of what other mechanisms activate these channels. An appealing hypothesis is that unknown ligands bind the PMD and the ligand-binding energy translates into movement of the linked S4 helix. Alternatively, interactions with the associated large polycystin-1 proteins, or to their unknown functions, may alter or initiate channel gating.

PI(4,5)P_2_ was recently observed to facilitate the gating of both polycystin-2 and polycystin 2-l1 by interacting with residues on both the N and C termini(Zheng et al., 2018). The current lack of resolution of N- and C-terminal ion channel flexible regions constrains our interpretation of gating influences for the group II TRP channels. Among group I TRP channels, the large interaction surfaces for intracellular ligands, such as the ankyrin repeats found on several TRPs, the more common TRP domains, and predicted C-terminal EF-hand calcium-binding sites and calmodulin-binding domains are difficult to interpret without better understanding of proteins or ligands that may bind these regions.

The explosion of structures available for TRP channels lays the groundwork for interpreting future detailed characterization. The structure of polycystin 2-l1 presented here, will help in understanding what drove evolution to create the diversity that exists in the TRPP subfamily, but only when we more completely understand their biophysical and physiological functions. The recent progress made in cryo-EM underscores the importance of discovering these roles.

## Methods

### Cloning, Expression and Purification

Cloning, expression and purification of the full length were completed using the BacMam strategy(Goehring et al., 2014). Briefly, constructs were subcloned into the vector pEG using restriction sites NotI and BstBI. The construct sequence was verified and transformed into DH10bac cells. Blue-white screening facilitated colony selection to create a midiprep of the bacmid. This preparation was used to transfect Sf9 cells using Cellfectin II following the manufacturer’s instructions. Amplification of the virus was completed for 2 cycles (P2) before viral particle were used to infect HEK293S GnTl-cells at 10% v/v at 37°C and 8% CO_2_ with shaking. After 24h, sodium butyrate was added to a final concentration of 10mM and cells were harvested after 48h.

Protein was extracted directly from pellets using 40x critical micellar concentration (CMC) of the detergent C12E9 in HKN buffer (in mM: HEPES 50, KCl 150, NaCl 50, CaCl_2_ 5; pH 7.5) with an EDTA-free protease inhibitor cocktail (Roche) tablet for 2h at 4°C using gentle rotation. Ultracentrifugation of the sample was completed at 40,000 RPM for 1h at 4°C. The resulting supernatant was retained and used for a batch incubation with amylose resin overnight at 4°C. Resin was collected, washed in an HKN buffer with higher K^+^ (500mM) buffer with 2x CMC of *n*-Dodecyl β-D-maltoside: cholesteryl hemisuccinate for 10 bed volumes followed by a normal HKN buffer for 10 bed volumes with 2x CMC of DDM: CHS. The protein was eluted using 40 mM maltose in HKN buffer at 2x CMC with DDM: CHS. All fractions were collected, concentrated and subjected to 1 round of size exclusion chromatography using an Increase Superose6 (GE Lifesciences) column pre-equilibrated with HKN and 2xCMC DDH: CHS. Protein-containing fractions, as determined by A280 and western-blotting against maltose binding protein (MBP), were collected, concentrated and used for further preparation. The sample’s concentration was determined using A280 and incubated with a 1:3 ratio (mass) of poly (maleic andydride-alt-1-decene substituted with 3-(dimethylamino) propylamine; PMAL C8) overnight with gentle rotation at 4°C. A final round of size exclusion chromatography using an Increase Superose6 column equilibrated with HKN buffer was performed, fractions collected, and concentrated using a Vivaspin Turbo4 100,000 molecular weight cutoff (MWCO) centrifugation device before being applied to grids and freezing.

### Sample Preparation

The cryo specimen was vitrified using 3ul of purified polycystin 2-l1 in PMAL-C8 at 3.5mg/ml and applied onto a glow-discharged 400 mesh copper Quantifoil R1.2/1.3 holey carbon grid (Quantifoil). Grids were blotted for 3s at 95% humidity and flash frozen in liquid nitrogen-cooled liquid ethane bath using a Vitrobot (Thermo Fisher, Hillsboro OR). Samples were stored in liquid nitrogen until acquisition.

### CryoEM Image Acquisition, Processing and Modeling

Data for full-length polycystin 2-l1 were collected on a FEI Titan Krios (Thermo Fisher, Hillsboro OR) at 300kV with a Gatan Quantum Image Filter (20 eV slit) and K2 Summit detector at the Janelia Research Campus (Ashburn, VA) using the following parameters: A total of 3,814 image stacks were acquired at a sampling rate of 1.04Å/pixel with an 8s exposure of 8e- pixel^−1^ s^−1^ at 0.2s/frame for a total dose of approximately 60e-Å^2^ using SerialEM(Mastronarde, 2005).

Data was processed using cisTEM(Grant et al., 2018), which contains all processing steps listed below with relevant references to the technique for each step. Briefly, beam-induced motion and physical drift were corrected followed by dose-weighing using the Unblur algorithm (Grant and Grigorieff, 2015). Next the contrast transfer function was estimated and used to correct micrographs. After inspection of all micrographs, 3,494 were used for data processing. Particles were then automatically selected based on an empirical evaluation of maximum particle radius (80Å), characteristic particle radius (60Å), and threshold peak height (2 S.D. above noise). 2D classification on 842,130 particles employed an input starting reference of a previous model solved in C4 symmetry (Supplemental Figure 2). This initial model was generated *ab initio* from a data set acquired for polycystin 2-l1 and processed using cisTEM (particles from 20Å to 8Å for the first step of classification with a 20 Å low pass filter used for the initial model to avoid bias). Eight class averages representing different orientations were selected for further iterative 3D classification and refinement in C4 symmetry (Supplemental Figure 1). The best solutions for each iteration were selected for local refinement in (Supplemental Figure 2) cisTEM. Upon completion of 3D refinement, the best class was sharpened using B-factors. FSC and angular distribution plots were collected for the final dataset in cisTEM (Supplemental Figure 3).

Polyalanine *α*-helices and *β*-strands were built for each transmembrane section and the PMD using pdb 5T4D as a final guide in Coot(Emsley et al., 2010). Densities were inspected, and initial assignments of residues were based on aromatic or bulky residues. Real-space refinement, using PHENIX(Adams et al., 2010), was completed iteratively with inspection of the model in Coot. Model quality was evaluated using Molprobity(Williams et al., 2018) and EMRinger(Barad et al., 2015). The model was then compared to sequence alignments of the TRPP family and three models of the closely related polycystin-2 structures(Grieben et al., 2017, Shen et al., 2016, Wilkes et al., 2017). Pore size was evaluated using HOLE(Smart et al., 1996) and local resolution maps calculated from RESMAP(Kucukelbir et al., 2014). The sequence analysis for polycystin 2-l1 family members and display onto the structure was completed using 500 multiple sequence alignments generated from iterative Blast searches and then analyzed with Consurf(Glaser et al., 2003, Landau et al., 2005). Electrostatic plots were created in Pymol. Model rendering was completed in Pymol and Chimera(Pettersen et al., 2004).

### Electrophysiology

Liposomes containing full-length and truncated polycystin 2-l1 protein were voltage clamped to record single channel currents. Protein was reconstituted following the dilution method (Cortes and Perozo, 1997) into liposomes made from soy extract polar (Avanti). After extensive dialysis, lipids were pelleted and resuspended in HKN buffer and stored at −80°C until characterization. A sample of reconstituted protein was dried overnight at 4°C under constant vacuum and rehydrated the next morning in the same buffer. Liposomes were allowed to swell for 1h on ice before recording with patch pipettes (10-15 MΩ). The bath was HKN buffer, pH 7.4; the pipette contained HKN buffer with 1mM MgCl_2_. Solution osmolarity was 390-400mOsm. Data was acquired at 10kHz and a low-pass Bessel-filtered at 1-2kHz. For each recording, pipette offset and capacitance were corrected. Approximately 2 min after a GΩ seal was achieved, the patch was excised, and capacitance corrected before recording.

**TABLE I.**
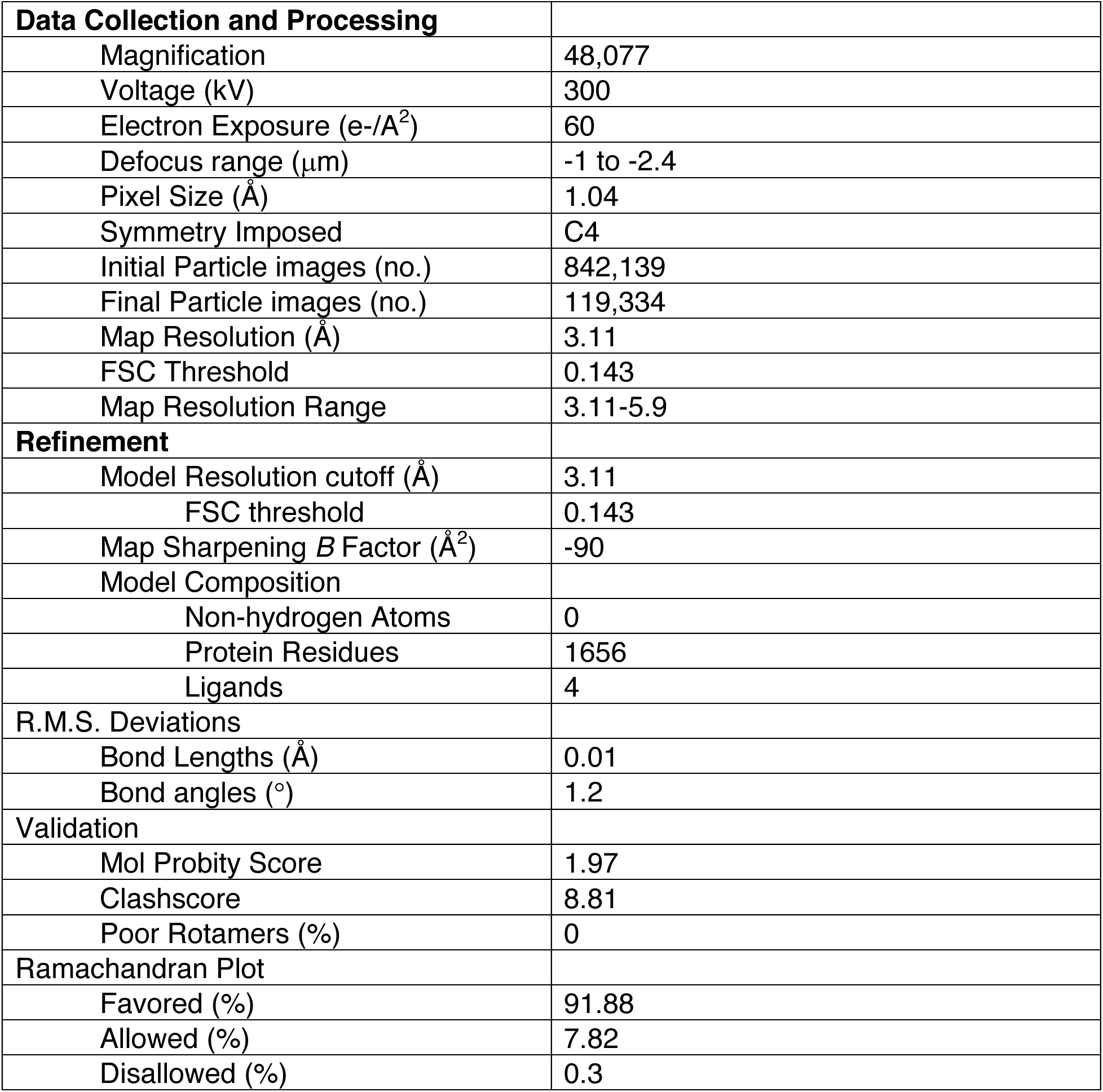
Cryo-EM data collection, refinement, and validation

## SUPPLEMENTARY FIGURES

**Supplemental Figure 1.**
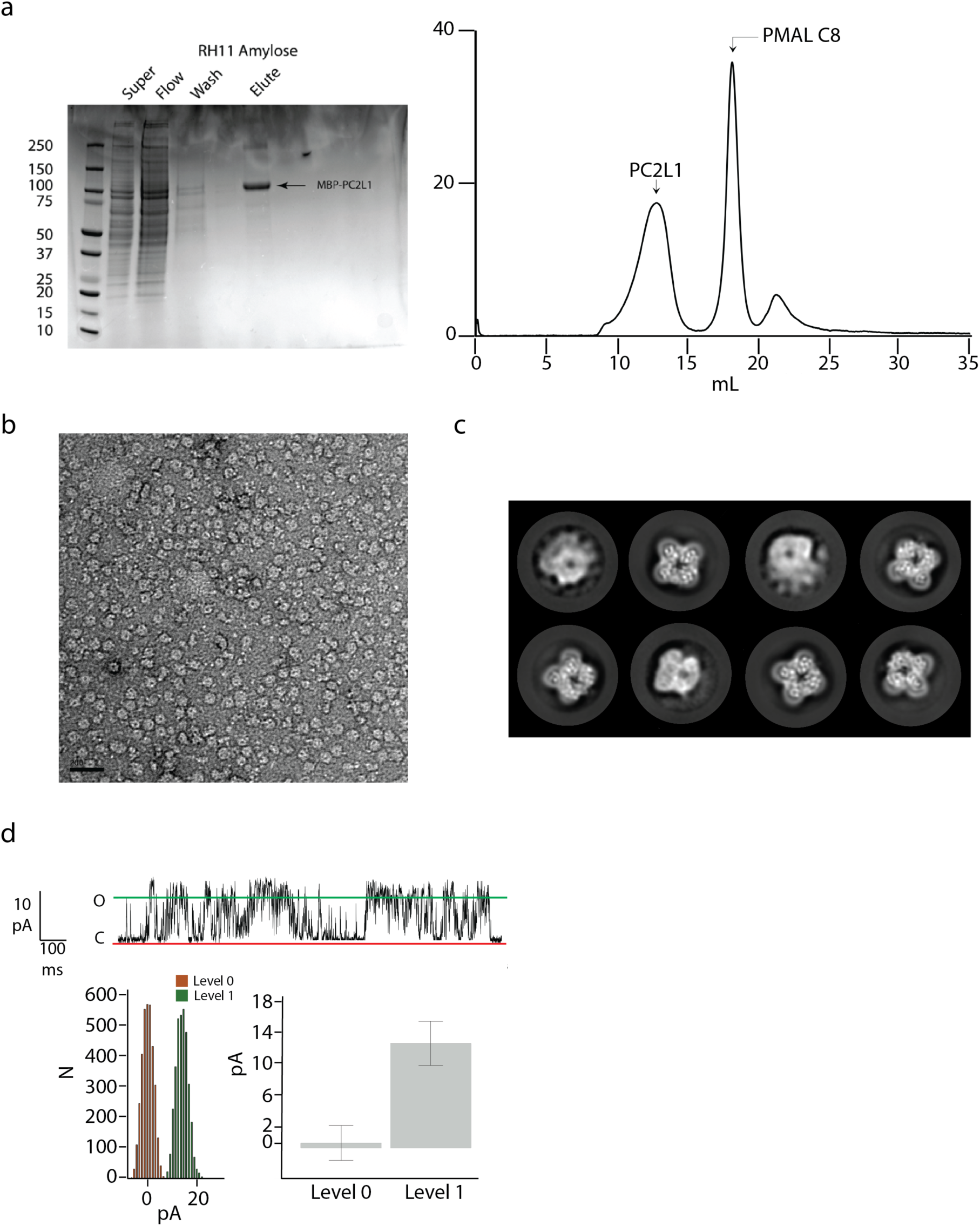
Purification, negative staining, 2D class average and function of polycystin 2-l1 a) SDS-PAGE of amylose purification for supernatant, flow, wash and elution (left) and size exclusion chromatography (Superose 6 Increase) after affinity chromatography and exchange from detergent to amphipol PMAL-C8 (right). b) representative sample of negative-stained polycystin 2-l1 in PMALC8 (scale bar, 200 nm). c) 8 representative 2D class averages used in data processing. (d) Sample trace of voltage-clamped, liposome-reconstituted recombinant polycystin 2-l1 (top). Amplitude histogram of open and closed events (bottom left) and mean current of open (Level 1) and closed (Level 0) events. Average current = 13.2pA (± 2.8 S.D.) at + 125mV, 1.6s duration.

**Supplemental Figure 2.**
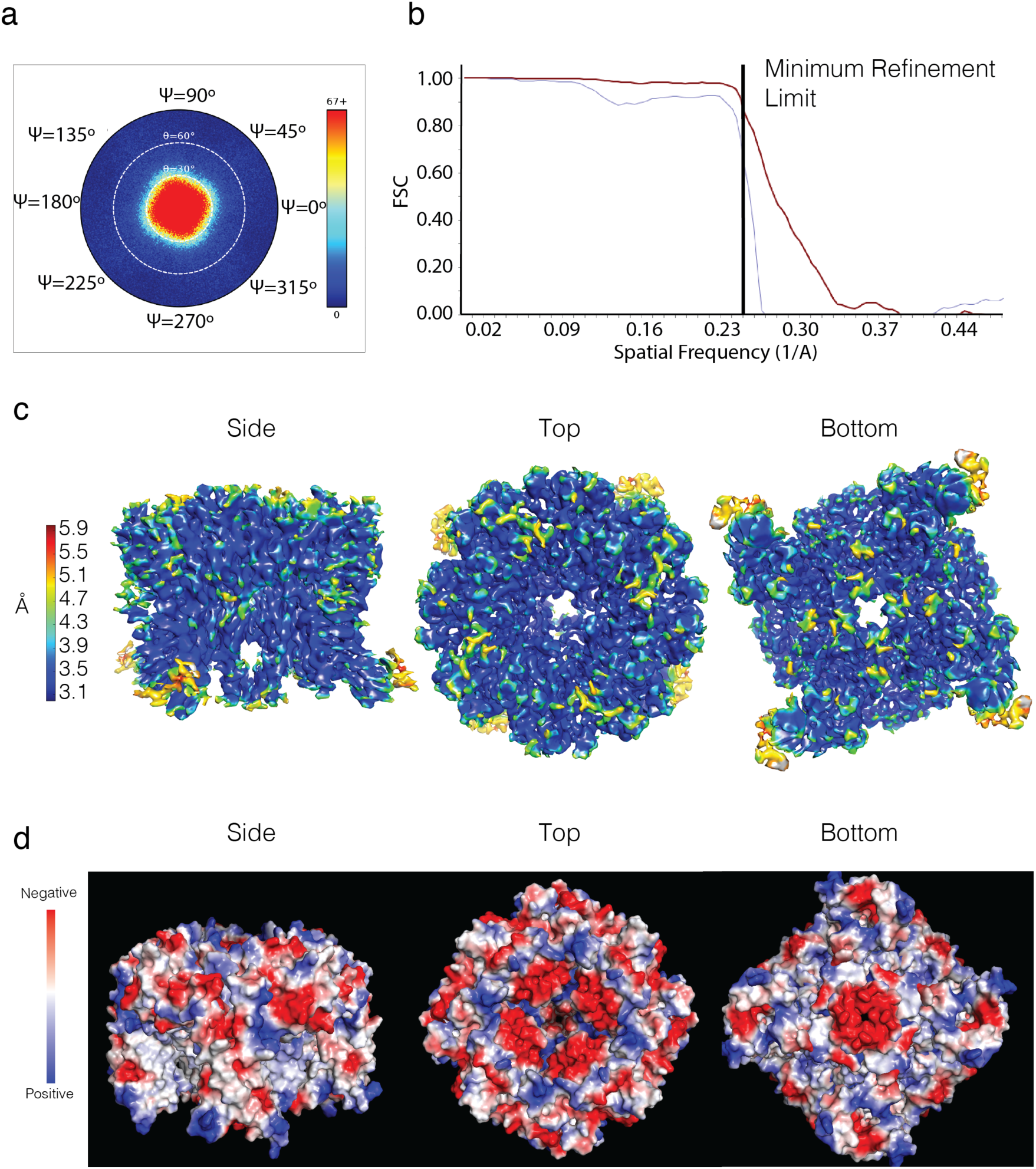
Fourier Shell Correlation (FSC) curve, local resolution and electrostatic plots of polycystin 2-l1 a) angular distribution plot and b) corresponding FSC of polycystin 2-l1 in PMAL C8 c) local resolution map determined by RESMAP d) electrostatic potential of polycystin 2-l1 in PMAL-C8.

**Supplemental Figure 3.**
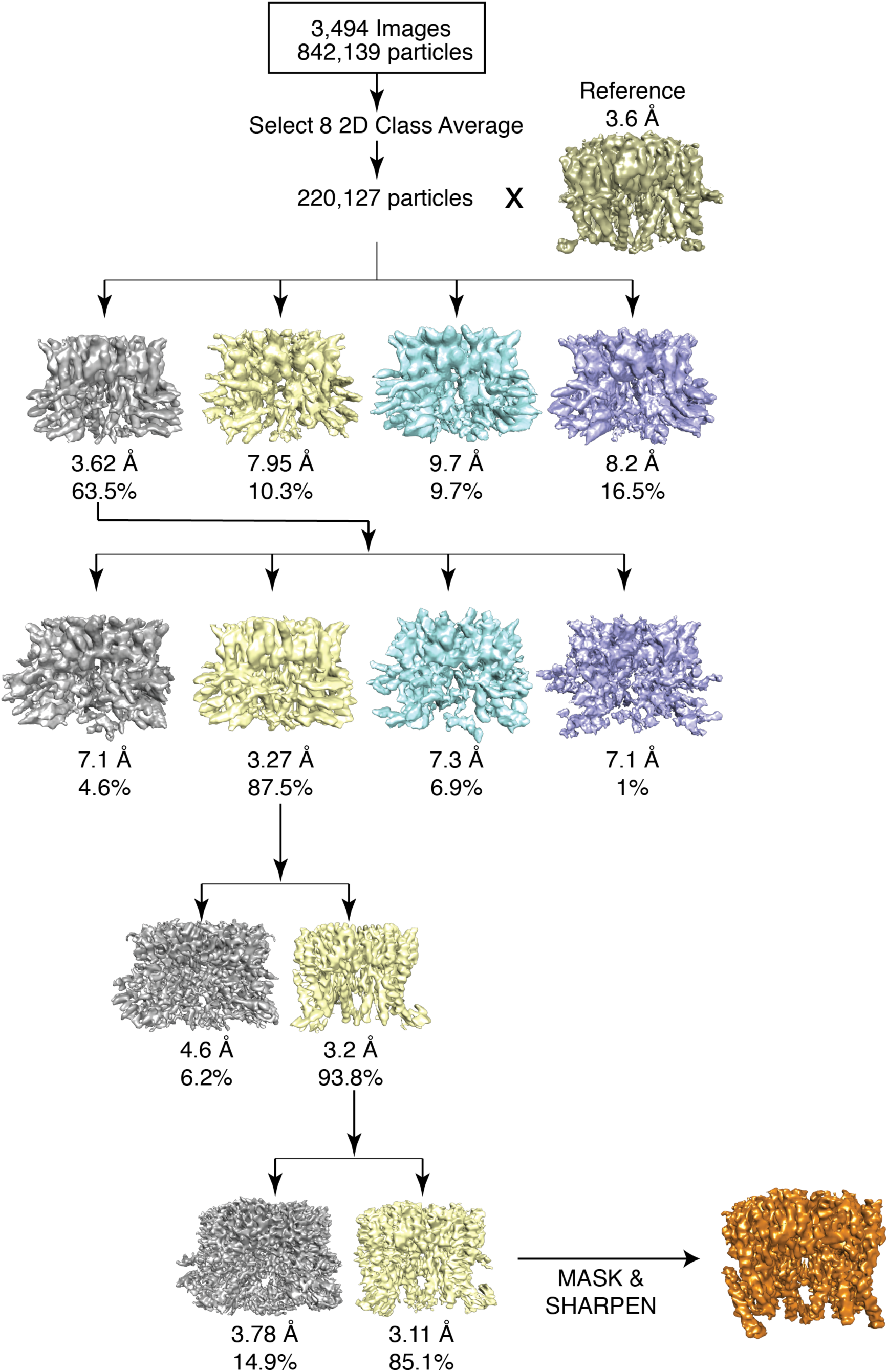
Single particle data processing. 842,139 particles were automatically selected to generate a 2D class average. Eight classes were selected; a reference from a previous dataset that generated a 3.6 Å map was used to derive four initial classes. The four classes were generated from automatic refinement; one class was selected to generate an additional round of four classes. The subsequent particle was then selected to generate two more classes (two more rounds with a mask to omit contributions from the amphipol). The final, highest resolution structure was selected and then sharpened.

## Acknowledgements

The authors would like to extend their gratitude to members of the Clapham, Grigorieff labs and the staff of the Cryo-EM facility at Janelia Research Campus for their feedback and support.

